# Compounds toxic to stationary-phase yeast reveal a role for lipid metabolism in quiescent cell survival

**DOI:** 10.64898/2026.06.13.732029

**Authors:** Minakhi Halder, Akira Honda, Yuki Kiyota, Koichiro Awai, Ayumu Yamamoto

## Abstract

The mechanisms that enable quiescent cells to survive nutrient-limited conditions remain imcompletely understood. To investigate these mechanisms, we performed a large-scale chemical screen to identify compounds that reduce viability of stationary-phase fission yeast cells. We identified two structurally related compounds that impair quiescent cell survival. In stationary phase, these compounds inhibited cell cycle re-entry, altered lipid droplets (LDs), and affected nuclear and chromosomal size, all of which are associated with lipid metabolism. In proliferating cells, they induced cell cycle arrest accompanied by nucleolar accumulation of cyclin-dependent kinase (CDK), a hallmark of stationary-phase cells, and inhibited mitochondrial oxygen consumption. The lipid synthesis inhibitor cerulenin similarly inhibited cell cycle re-entry and induced cell cycle arrest with nucleolar CDK accumulation; however, it did not reduce viability in stationary-phase cells or reproduce the other cellular effects. These results indicate that the compounds may affect lipid metabolism potentially by impairing lipid mobilization. Furthermore, the compounds were also toxic to stationary-phase budding yeast, suggesting conservation of survival mechanisms in quiescent cells. Together, our findings suggest that lipid metabolism, possibly lipid mobilization, plays a critical role in quiescent cell survival.

## INTRODUCTION

Upon nutrient starvation, cells cease cell division and enter a state called quiescence, which is characterized by reduced metabolic activity and resistance to environmental stresses. A growing body of evidence has shown that a subpopulation of pathogenic microorganisms is in quiescence [1–5]. These quiescent pathogenic organisms exhibit resistance to antimicrobial drugs, posing a major obstacle for treating infectious diseases [1,3,5,6]. Therefore, understanding the survival mechanisms of quiescent cells is of clinical importance.

Stationary phase is a quiescent state that microorganisms enter when carbon sources are exhausted. Studies in yeast have demonstrated that stationary-phase cells differ significantly from exponentially proliferating cells (log-phase cells) [6–12]. Stationary-phase cells exhibit reduced metabolic activity, enhanced stress resistance, and undergo extensive structural remodeling, including changes in cell and nuclear size and reorganization of cellular structures [13–29].

Previous studies of the fission yeast, *Schizosaccharomyces pombe* have shown that cells actively establish stationary phase in nutrient-poor, stressful environments. Although cyclin-dependent kinase (CDK), which drives the cell cycle, is inhibited during stationary phase [7,30–33], CDK also plays an active role in stationary-phase establishment [13]. Additionally, CDK accumulates at the nucleolus in stationary-phase, and this nucleolar accumulation has been suggested to be part of the stress response, as heat stress also induces it [13].

Moreover, lipid metabolism plays a crucial role in stationary phase. In stationary phase, lipid droplets (LDs) that store neutral lipids increase significantly [34–38]. A decrease in lipids or impaired lipid mobilization results in reduced ATP levels and decreased viability in cells in stationary phase or under glucose starvation [39,40]. In addition, defective lipid mobilization leads to significant delays in cell cycle resumption, indicating that stationary-phase cells rely on intracellular lipids for re-start of cell cycle [41,42].

Here, to understand the survival mechanisms of quiescent cells, we screened a chemical compound library for compounds that impair survival of *S. pombe* cells in stationary phase. We identified two structurally related compounds that affect processes associated with lipid metabolism. Their effects partially resemble those of the lipid synthesis inhibitor cerulenin, suggesting that the compounds affect lipid metabolism. Our findings suggest that lipid metabolism plays an important role in quiescent cell survival.

## MATERIALS AND METHODS

### Yeast strains, media, and culturing methods

For analyses of *S. pombe* cells, the following strains were used: L972 (*h^-^*), SMG25-1A (*h^+^ lys1^+^*::*ccr1N-NeonGreen cdc2^+^*::*GFP-kan^r^ hta1^+^*::*mCherry-nat^r^*), SMH2-9B (*h^+^cdc2^+^*::*GFP-kan^r^ hta1^+^*::*mCherry-nat^r^*), SMHT16-1 (*h^+^ lys1^+^*::*ccr1N-GFP hta1^+^*::*mCherry-nat^r^*), and SRS6-2D (*h^−^ cyc1*:: *kan^r^*). YES medium and basic genetic techniques were described previously [43]. Log-phase or stationary-phase cells were prepared by culturing cells to a density of approximately 5 × 10⁶ or 1.2 × 10⁸ cells/mL, respectively, with cell growth monitored every 2 hours. To assess cell cycle re-entry, stationary-phase cells were resuspended in YES medium without glucose (YES–Glc) and treated with compounds for 24 hours. Subsequently, an equal volume of YES medium containing 6% (w/v) glucose was added to the culture. For compound treatments, a DMSO stock solution of each compound was added to the culture at 1/100 of the final volume, and compound-free DMSO was used as a control.

For viability assays of *S. cerevisiae,* strain W303a (*MATa ade2-1 his3-11 leu2-3 trp1-1 ura3-1 can1-100*) was used. Stationary-phase *S. cerevisiae* cells were prepared by culturing cells in YPAD medium (1% yeast extract, 2% peptone, 2% glucose, and 40 mg/L adenine sulfate) at 30 °C until the density reached ∼3 × 10⁸ cells/mL, followed by an additional 7-day incubation. Stationary-phase cells were treated with compounds after the culture was adjusted to a density of 5 × 10^6^ cells/mL.

### Viability assay

Viability was assessed by colony formation. Compounds were removed by washing cells with medium prior to plating on YES solid medium. A total of 110 individual cells were placed on YES (for *S. pombe*) or YPAD (for *S. cerevisiae*) solid medium using a fine microneedle under a microscope and incubated at 32 or 30°C, respectively.

### Screening method of the compounds

A library of chemical compounds used for screening was generated by Takeda Pharmaceutical Company Limited. In the first screening, after cells reached stationary phase in YES medium, the culture medium was replaced with YES–Glc, and the cells were incubated overnight at 30°C. The culture was then diluted to a concentration of 5 × 10⁶ cells/mL, and 100 µL of the diluted culture was incubated with each compound in 384-well plates at 30°C for 24 hours. Subsequently, glucose was replenished by adding 100 µL of YES medium containing 6% (w/v) glucose to each well, and cultures were incubated for another 24 hours at 30°C. Cell proliferation was assessed by measuring OD_600_, and compounds that inhibited cell proliferation were selected. In the second screening, 100 µL of log-phase culture was incubated with each selected compound at 30°C in 384-well plates. Cell growth was monitored for 9 hours by measuring OD_600_, and compounds that did not inhibit cell proliferation were selected.

### Visualization of cell, nuclear membrane, chromosomes, and Cdc2

The nuclear membrane was visualized using a gene encoding the GFP-tagged N-terminal portion of the NADPH-cytochrome P450 reductase (Ccr1N-GFP) [44], or NeonGreen-tagged Ccr1N (see below). Chromosomes were visualized using a gene encoding an Hta1-mCherry fusion protein [45]. Cdc2 was visualized using the endogenous *cdc2^+^* gene fused to the *GFP* gene [13].

A plasmid carrying the *ccr1N-NeonGreen* gene was constructed based on an integration plasmid pYK3, generated by inserting restriction sites at the beginning of the Ccr1N coding region in pMH3 harboring the *ccr1N-GFP* gene [44]. A DNA fragment encoding NeonGreen was amplified by PCR using pFA6a-link-ymNeongreen-SpHis5 (a gift from Bas Teusink, Addgene plasmid #125704; http://n2t.net/addgene:125704; RRID:Addgene_125704) as a template. The *GFP* gene on pYK3 was replaced with the *NeonGreen* gene by homologous recombination in bacterial cells [46], using the PCR product. The resulting *ccr1-NeonGreen* plasmid, pMG4, was introduced into cells for nuclear membrane visualization.

### Quantification of cell and nuclear volume, the chromosome-occupying area, and nucleolar accumulation of Cdc2

Images were acquired and analyzed, as previously described [13]. Log-phase and stationary-phase cells were suspended in EMM and EMM without glucose (EMM–Glc) medium, respectively. When required, compounds were added to the medium. Approximately 20 µL of cell suspension was placed on a 40 × 50 mm coverslip (Matsunami Glass Ind., Ltd., Osaka, Japan) coated with 5 mg/mL lectin (Sigma-Aldrich Japan, Tokyo). Observations were made using an IX71 inverted microscope equipped with a CoolSNAP-HQ2 CCD camera (Nippon Roper Co. Ltd., Tokyo) and a 60 × /1.42 NA Plan Apo oil-immersion objective lens (Olympus Corp., Tokyo). Images of the nuclear envelope, chromosomes, and Cdc2 were acquired at 11 to 15 focal planes spaced at 0.3 μm intervals. Images were deconvolved and analyzed using MetaMorph (version 7, Molecular Devices Japan) and ImageJ [47]. Cell volume was calculated using the Pombe Measurer ImageJ plugin (http://www.columbia.edu/~zz2181/Pombe_Measurer.html). Nuclear area was determined by fitting an ellipse to the clearest section of the nuclear envelope. Nuclear volume was calculated assuming the nucleus is a spheroid with semi-axes lengths, a, b, and (a+b)/2. Chromosome area was measured from binarized 2D projections of a deconvolved set of images. The nucleolar accumulation index (NAI) of Cdc2 was determined as the ratio of Cdc2 signal intensity in the nucleolar region to the chromosomal region, using 2D projections generated by additive image projecting method, as described previously [13].

### Quantification of intracellular LD signal intensity

Cells were cultured to stationary phase in YES medium and transferred to YES–Glc medium at a density of ∼5 × 10⁶ cells/mL. Cells were then incubated at 30°C with compounds or DMSO (control) for 5 days. LDs were visualized as follows. Cells in 250 µL culture were resuspended in an equal volume of fresh YES–Glc medium. Cells were stained by adding 1 µL of 250 µM BODIPY in DMSO and incubating for 10 minutes at room temperature. The stained cells were resuspended in 2 µL PBS, and a 1.5 µL aliquot was placed on a 24 × 60 mm slide glass and covered with an 18 × 18 mm coverslip. LD images were acquired and processed as described for nuclei, chromosomes, and Cdc2. Cell area was defined by polygonal selection using brightfield images. The LD signal intensity within the cell area was measured using 2D projections generated by additive image projection method, with background intensity subtracted from the total fluorescence intensity.

### Measurement of oxygen consumption rate

Oxygen consumption in log- or stationary-phase cells was measured at 25°C using a Clark-type oxygen electrode (Hansatech Instruments Ltd.). For each assay, 2 mL of cell suspension containing 1 × 10⁷ cells/mL was used.

## RESULTS

### Screening of chemical compounds toxic to stationary-phase cells

To identify compounds that target survival of *S. pombe* cells in stationary phase, we screened a chemical library for compounds that were toxic to stationary-phase cells but not to log-phase cells by a two-step screening approach (Fig 1A and 1B). From approximately 60,000 compounds, 320 were identified in the first screening and 18 in the second screening. We then re-evaluated these 18 compounds using standard culture procedures. Eventually, two structurally related compounds, designated 0511 and 0512, consistently inhibited cell cycle re-entry of stationary-phase cells without affecting multiple rounds of cell division of log-phase cells (Fig 1C).

**Fig 1.**
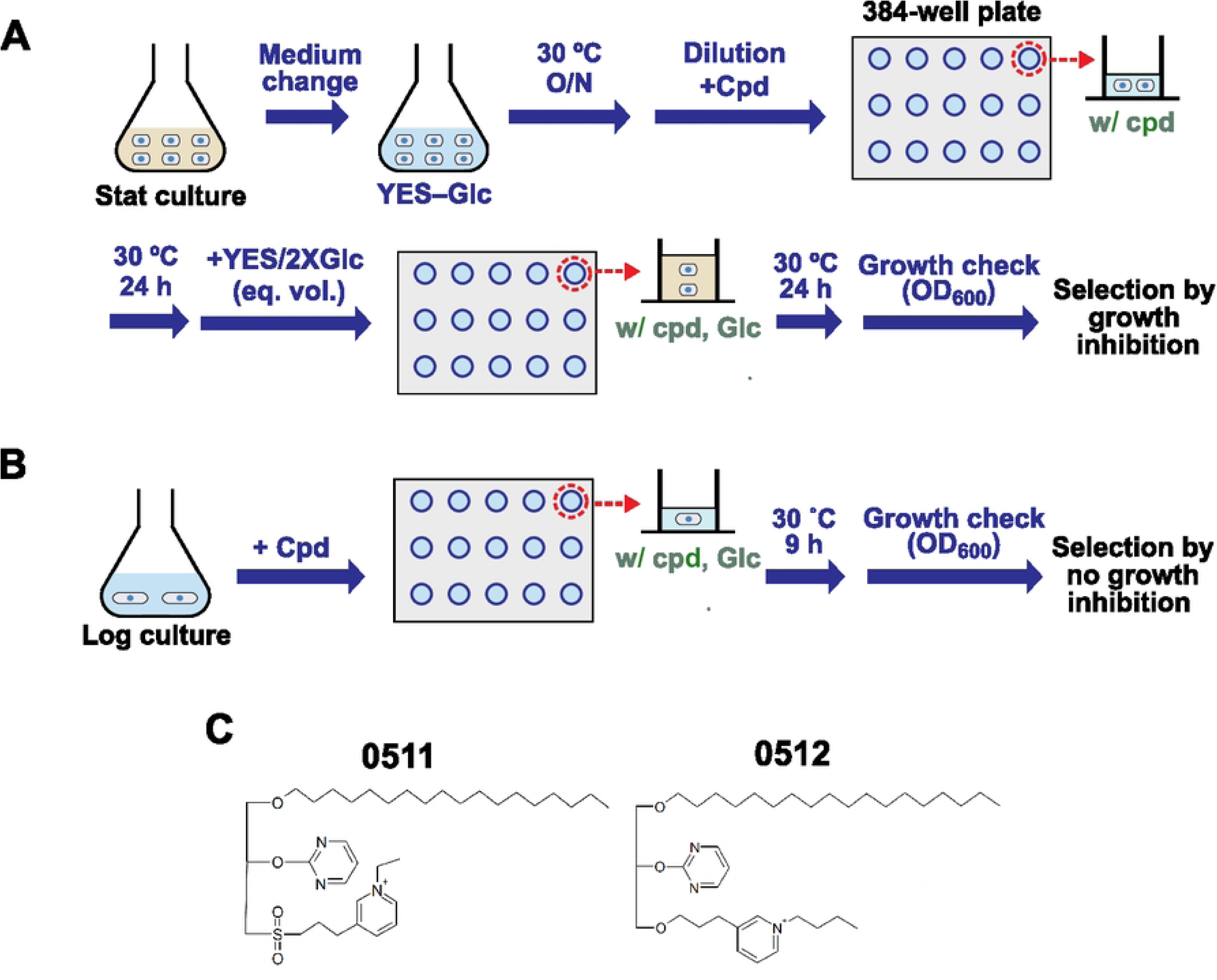
Screening of chemical compounds toxic to stationary-phase cells. (A) First screening for compounds that inhibit cell cycle re-entry in stationary phase. Stationary-phase cells grown in YES rich medium (Stat culture) were transferred to fresh YES medium without glucose (YES–Glc), which has previously been shown to extend survival of stationary-phase cells [13]. After overnight incubation at 30 °C, the culture was transferred to wells of a 384-well plate, and various chemical compounds (Cpd) were added. After 24 hours of incubation at 30 °C, an equal volume of YES medium containing twice the glucose concentration (YES/2 × Glc) was added to each well to replenish glucose. Following an additional 24 hour incubation at 30 °C, cell proliferation in each well was assessed by measuring OD_600_. (B) Second screening for compounds that do not inhibit proliferation of log-phase cells. The culture containing log-phase cells (Log culture) were transferred to wells of a 384-well plate along with the chemical compounds selected from the first screening. After 9 hours of incubation at 30°C, cell proliferation was monitored by OD_600_. (C) Chemical structures of the identified compounds. Strain used: L972.

### 0511 and 0512 inhibit cell cycle re-entry and reduce viability of stationary-phase cells

We examined the effects of 0511 and 0512 on stationary-phase cells in detail. Stationary-phase cells grown in YES medium were transferred to glucose-free YES medium (YES–Glc), which has previously been shown to extend survival of stationary-phase cells [13], and treated with each compound for one day (Fig 2A). Cell cycle re-entry was induced by adding an equal volume of YES medium with twice the glucose concentration. Both 0511 and 0512 inhibited cell cycle re-entry at 30 µM (Fig 2B), whereas neither compound affected re-entry at 10 µM (Fig 2C). We also found that inhibition of cell cycle re-entry requires the continued presence of the compounds. Removal of either compound restored cell cycle re-entry (Fig 2B). Although both 0511 and 0512 similarly inhibited cell cycle re-entry, their effects differed. 0511 allowed approximately one round of cell division, as indicated by a two-fold increase in cell density, and removal of 0511 rapidly restored re-entry. In contrast, 0512 completely inhibited cell cycle re-entry, and cells did not resume the cell cycle for at least 10 hours after its removal.

**Fig 2.**
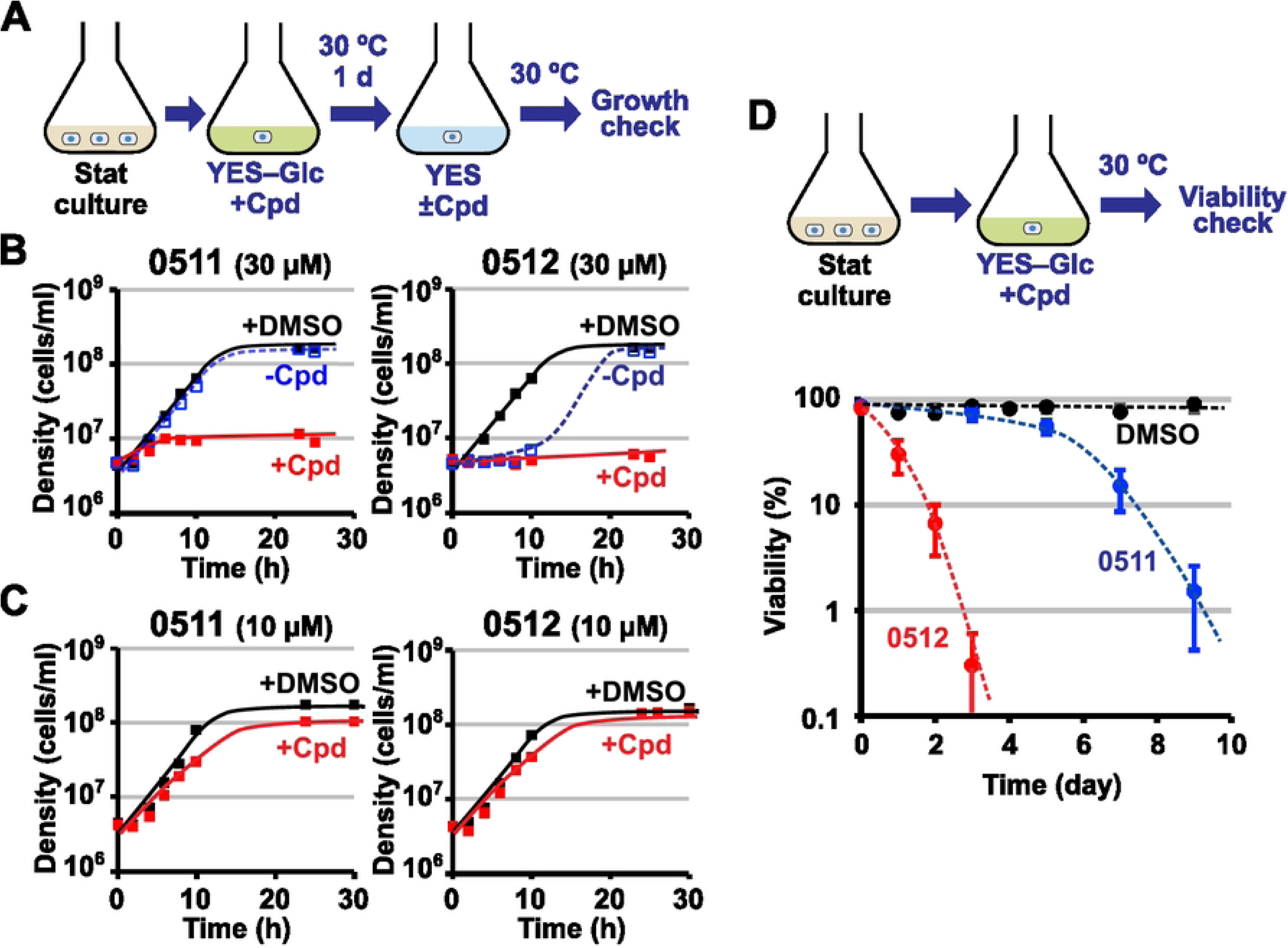
Effects of 0511 and 0512 on cell cycle re-entry and viability of stationary-phase cells. (A) Experimental design for assessing effects of 0511 and 0512 on cell cycle re-entry. Stationary-phase cells were treated with 0511 or 0512 in YES–Glc medium for 1 day at 30 °C and then tested for their ability to re-enter the cell cycle in YES medium, both in the presence and the absence of the chemical compounds (Cpd). (B and C) Effects of 0511 and 0512 on cell cycle re-entry of stationary-phase cells at 30 µM (B) or 10 µM (C). Stationary-phase cells treated with 0511 or 0512 in YES–Glc medium were examined for proliferation in YES medium. Growth curves of DMSO-treated cells in the presence of DMSO (+DMSO) and compound-treated cells in the presence (+Cpd) or absence of Cpd (–Cpd) of the compounds are shown. Each graph represents one of two biological replicates. (D) Effects of 0511 and 0512 on viability of stationary-phase cells. The upper schematic illustrates the culturing method used for viability analysis. The graph shows mean viability of stationary-phase cells treated with the compounds at 30 µM over time. Each mean value was calculated from three biological replicates. Error bars: standard error of mean (SEM). Strain used: L972.

Consistent with the screening strategy, colony formation assays showed that both 0511 and 0512 reduced cell viability, whereas the control DMSO caused no reduction for at least 10 days (Fig 2D). Viability declined more rapidly with 0512 than with 0511, which likely accounts for the delayed cell cycle re-entry observed for 0512-treated stationary phase cells (Fig 2B). These results show that both 0511 and 0512 reduce viability of stationary phase cells, with 0512 exhibiting greater toxicity.

### 0511 and 0512 inhibit the cell cycle progression and reduce viability of log-phase cells

We also examined the effects of 0511 and 0512 on log-phase cells in detail. When either compound was added at 15 or 30 µM (Fig 3A), cells initially proliferated like control DMSO-treated cells (Fig 3B and 3C), confirming the screening results. However, after approximately three cell divisions, cells ceased proliferation, unlike DMSO-treated cells. Treatment of log-phase cells with 0511 or 0512 at various initial cell densities consistently resulted in cell cycle arrest after about three cell divisions (Fig 3D). These findings suggest that the delayed cell cycle arrest was independent of nutrient levels or cell density and instead correlated with the number of cell divisions completed. Additionally, treatment with the compounds led to a more rapid decrease in cell viability compared to DMSO treatment (Fig 3E), indicating that both 0511 and 0512 also reduce viability of log-phase cells.

**Fig 3.**
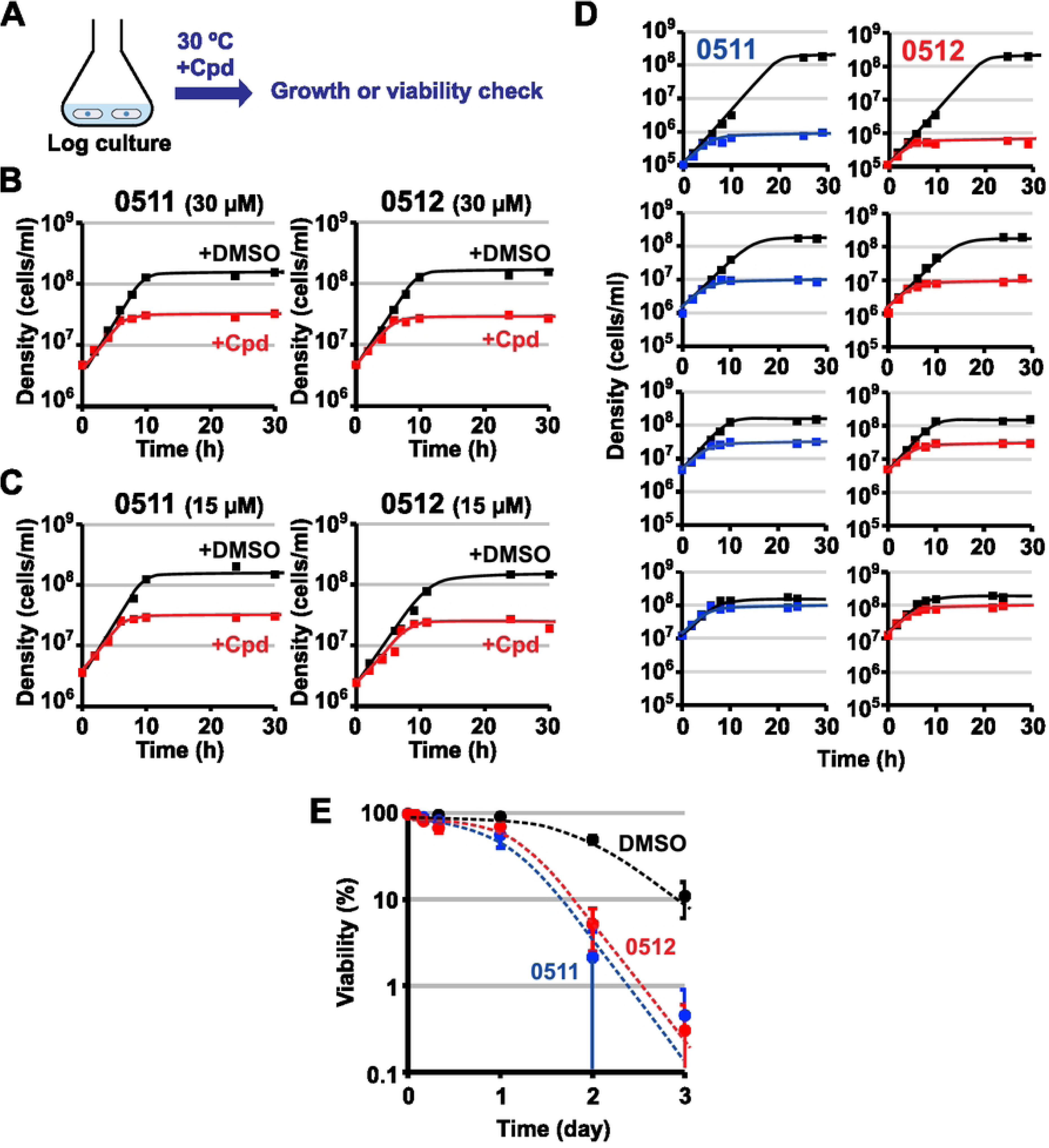
Effects of 0511 and 0512 on cell division and viability of log-phase cells. (A) Experimental design for assessing the effects of 0511 or 0512 on log-phase cells. (B and C) Effects of 0511 and 0512 on proliferation of log-phase cells at 30 µM (B) or 15 µM (C). Growth curves of log-phase cells treated with DMSO (+DMSO) or with 0511/0512 (+Cpd) are shown. Each graph represents one of two biological replicates. (D) Effects of 0511 or 0512 at 30 µM on proliferation at different cell densities. Compounds were added at time 0 h, and proliferation was monitored at various cell densities. (E) Effects of 0511 and 0512 on the viability of log-phase cells. The graph shows the mean viability of log-phase cells treated with the compounds at 30 µM over time. Each mean value was calculated from six biological replicates. Error bars: SEM. Strain used: L972.

### 0511 and 0512 induced accumulation of Cdc2 in the nucleolus upon cell cycle arrest

Since the compounds induce cell cycle arrest of log-phase cells, we investigated the effects of 0511 and 0512 on intracellular localization of Cdc2 (*S. pombe* CDK). In mono-nucleated log-phase cells, Cdc2 is uniformly distributed in the nucleus, whereas it accumulates in the nucleolus in stationary phase (Fig 4A) [13]. Similarly, when log-phase cells were treated with DMSO, Cdc2 is uniformly distributed in the nucleus (Fig 4B, Log and +DMSO, 8h) and localized in the nucleolus in stationary phase (Fig 4B, +DMSO, 24h). In contrast, treatment of log-phase cells with either compound induced nucleolar accumulation of Cdc2 at the time of cell cycle arrest, and the levels of nucleolar accumulation were comparable to those observed in stationary phase (Fig 4B and C; S1 Fig). These findings suggest that the compounds induce a stationary phase-like state. On the other hand, when stationary-phase cells were treated with 0511 or 0512, the nucleolar localization of Cdc2 remained unchanged, as seen in the control DMSO treatment (Fig 4D).

**Fig 4.**
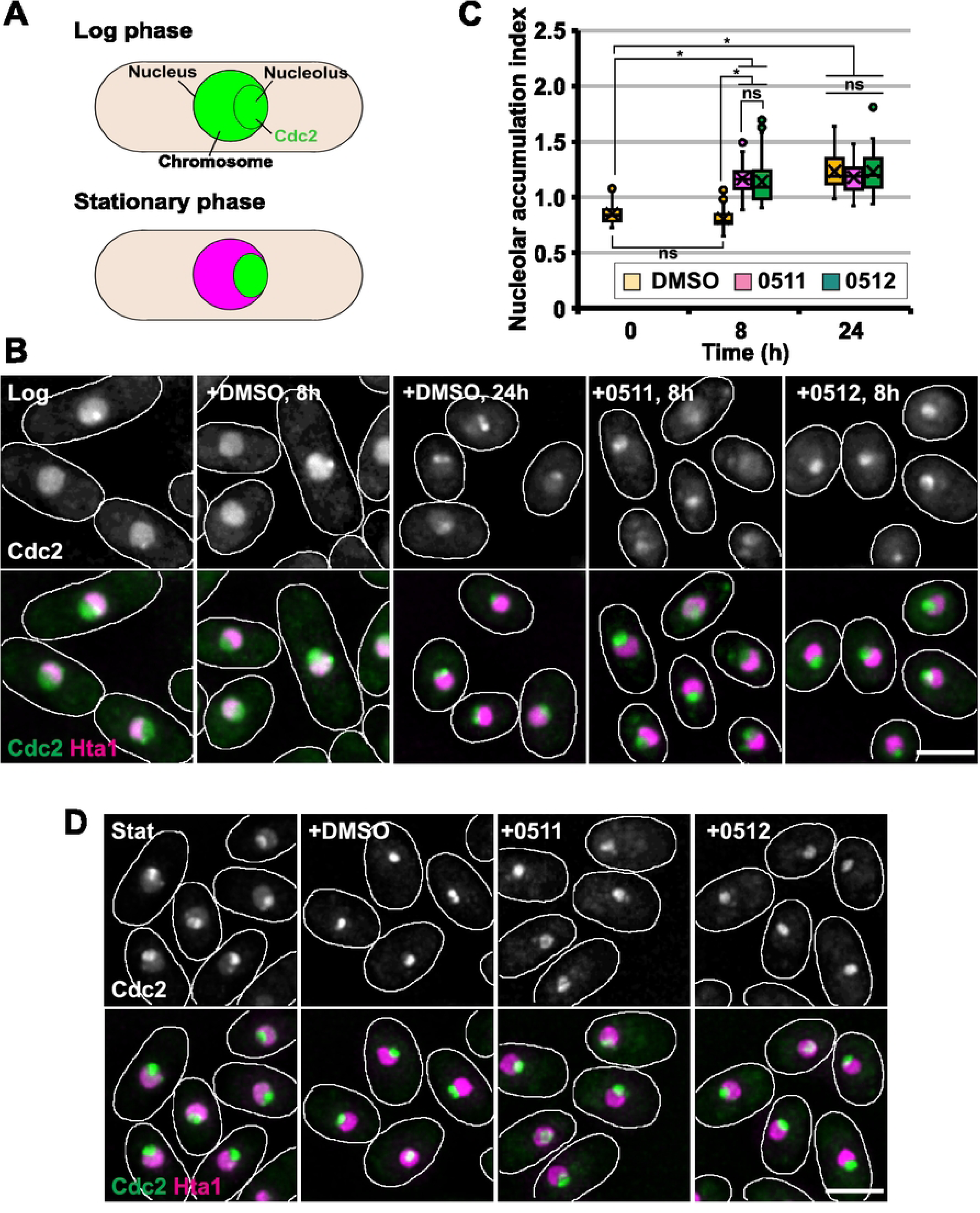
Effects of 0511 or 0512 on intracellular Cdc2 localization. (A) Schematic diagram of Cdc2 localization in log and stationary phase. (B) Cdc2 localization in cells treated with 0511 or 0512 during log phase. Log-phase cells producing GFP-tagged Cdc2 and mCherry-tagged histone H2A (Hta1) were treated with DMSO or the compounds as in Fig 3B or 3C. (C) Effects of 0511 and 0512 on Cdc2 signal intensity in the nucleolar region relative to the chromosomal region (nucleolar accumulation index). Cdc2 nucleolar accumulation was analyzed at the time of the cell cycle arrest (8 h) or after the arrest (24 h). More than 50 cells were analyzed at each time point. The horizontal line represents the median, the box indicates the interquartile range (IQR), whiskers show the remaining distribution within 1.5×IQR, circles outside whiskers represent outliers, and the cross indicates the mean value. (D) Cdc2 localization in cells treated with 0511 or 0512 during stationary phase. Stationary-phase cells were treated with DMSO or the compounds for one day. In (B) and (D), images show representative Cdc2 localization from one of two biological replicates, and white lines indicate cell outlines. Bar: 5 µm. Strains used: SMG25-1A (B and C); SMH2-9B (D).

### 0511 and 0512 affect nuclear and chromosomal size

Because 0511 and 0512 induced a stationary phase-like state, we examined the effects of the compounds on nuclear and chromosomal size, as well as on the nuclear-to-cell volume ratio (N/C ratio), all of which normally decrease during and after the establishment of stationary phase [13]. When log-phase cells were treated with DMSO, cell and nuclear volumes, the N/C ratio, and chromosomal size decreased during the transition to stationary phase, similar to the changes observed in non-treated cells (Fig 5A–5D, Treatment in log phase; S1 Fig). 0511 and 0512 initially allowed a slight decrease in cell size, but after cell cycle arrest, cell size was unchanged (Fig 5A, Treatment in log phase; S1 Fig). Furthermore, the compounds did not cause any significant decrease in nuclear and chromosomal size and increased the N/C ratio (Fig 5B–5D, Treatment in log phase; S1 Fig). These results indicate that although 0511 and 0512 induced stationary phase-like cell cycle arrest, they do not cause reductions in nuclear and chromosomal size, which are associated with stationary phase.

**Fig 5.**
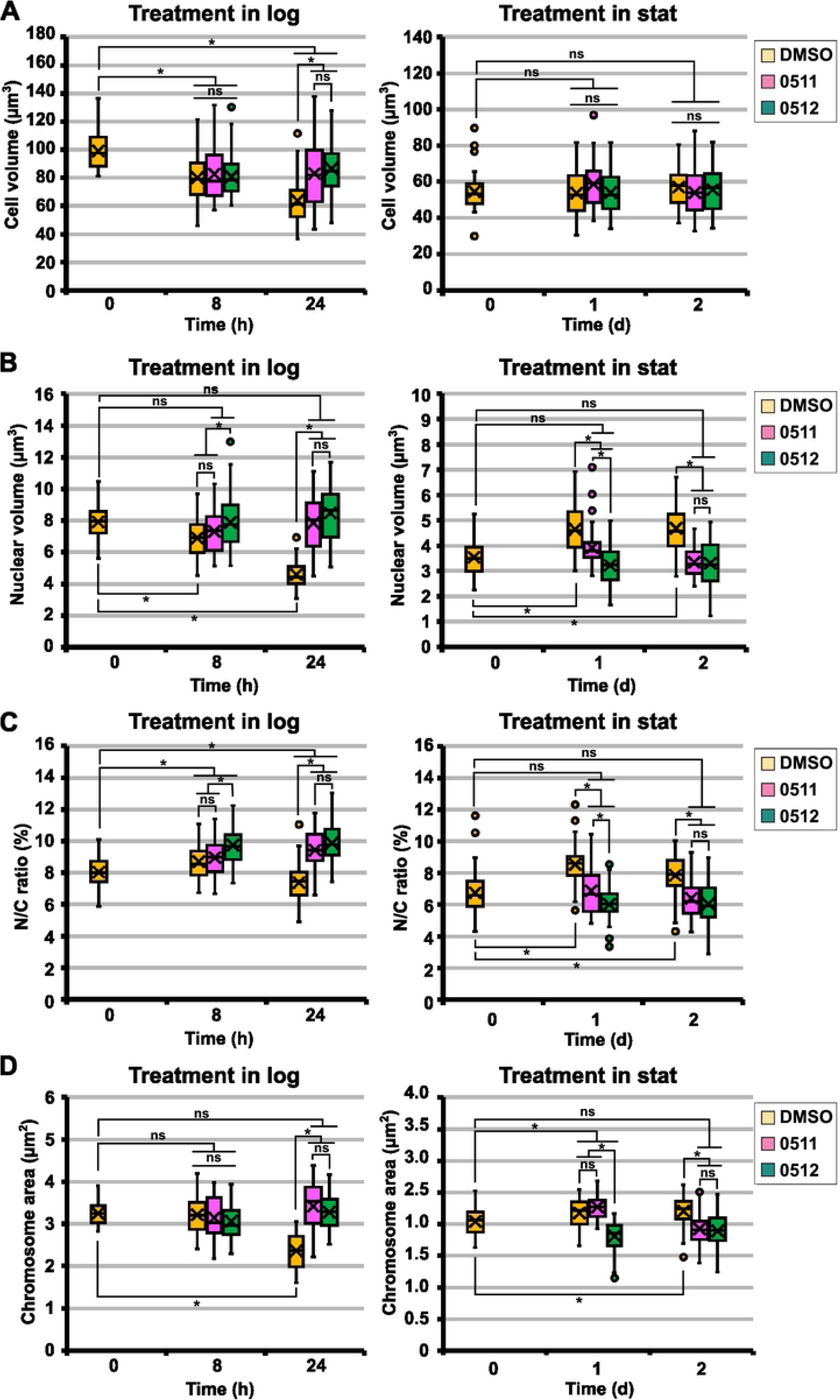
Effects of 0511 or 0512 on cell and nuclear size, and chromosome area. (A–D) The volumes of the cell (A) and the nucleus (B), the N/C ratio (C), and chromosome area (D) in cells treated with the compounds in log phase (Treatment in log) or in stationary phase (Treatment in stat). In compound-treated log-phase cells, size was analyzed at the time of cell cycle arrest (8 h) or after the arrest (24 h). Graphs show results from one of two biological replicates. More than 50 cells were analyzed at each time point. In all box-and-whisker plots, the horizontal line represents the median, the box indicates the interquartile range (IQR), whiskers show the remaining distribution within 1.5×IQR, circles outside whiskers represent outliers, and the cross indicates the mean value. Asterisks denote statistically significant differences (*P*<0.05). ns: not significant difference. Statistical analyses were performed using an unpaired, two-tailed Student’s *t-*test with the Bonferroni correction. Strain used: SMHT16-1.

The compounds also affected changes in nuclear and chromosomal size after stationary-phase entry. Nuclear and chromosomal size normally decrease even after entry into stationary phase [13]. However, control 1% DMSO treatment unexpectedly increased nuclear and chromosomal sizes and the N/C ratio without affecting cell size (Fig 5A–5D, Treatment in stationary phase; S2 Fig). In contrast, both 0511 and 0512 suppressed these DMSO-induced increases, resulting in nuclear and chromosomal sizes comparable to those of untreated stationary-phase cells after two-day treatment. All these findings suggest that the compounds affect the mechanisms regulating nuclear and chromosomal size during and after stationary-phase establishment.

### 0511 and 0512 affect LDs and inhibit mitochondrial oxygen consumption

Lipid metabolism plays a key role in cell cycle re-entry of stationary-phase cells and the regulation of nuclear size [41,42,48]. Given that 0511 and 0512 inhibit cell cycle re-entry and nuclear size reduction as well as their monoacylglycerol-like structures (Fig 1D), we hypothesized that these compounds might affect lipid metabolism. To explore this hypothesis, we examined the effects of 0511 and 0512 on lipid storage granules, LDs, using the lipid-specific dye BODIPY.

As previously reported [49], multiple punctate signals corresponding to LDs were observed by BODIPY staining in log-phase cells (Fig 6A, 0h). DMSO treatment for 4 hours did not alter the staining pattern or signal intensity, whereas 8-hour treatment led to a decrease in signal intensity (Fig 6A and 6B; S3A Fig), consistent with progression toward stationary phase, where BODIPY signals are reduced (Fig 6C and 6D). In contrast, 0511 treatment resulted in diffuse cytoplasmic staining along punctate signals, whereas 0512 treatment increased the size of puncta (Fig 6A). Both compounds significantly increased intracellular BODIPY signal intensity: 0511 caused a continuous increase up to 8 hours, whereas 0512 showed a slight decrease after 4 hours (Fig 6B; S3A Fig). In stationary-phase cells, neither 0511 nor 0512 altered punctate staining patterns, but both markedly increased intracellular BODIPY signal intensity compared to DMSO treatment (Fig 6C and 6D; S3B Fig). Collectively, the changes in staining patterns and signal intensities are consistent with an effect of 0511 and 0512 on lipid metabolism in both log-and stationary-phase cells.

**Fig 6.**
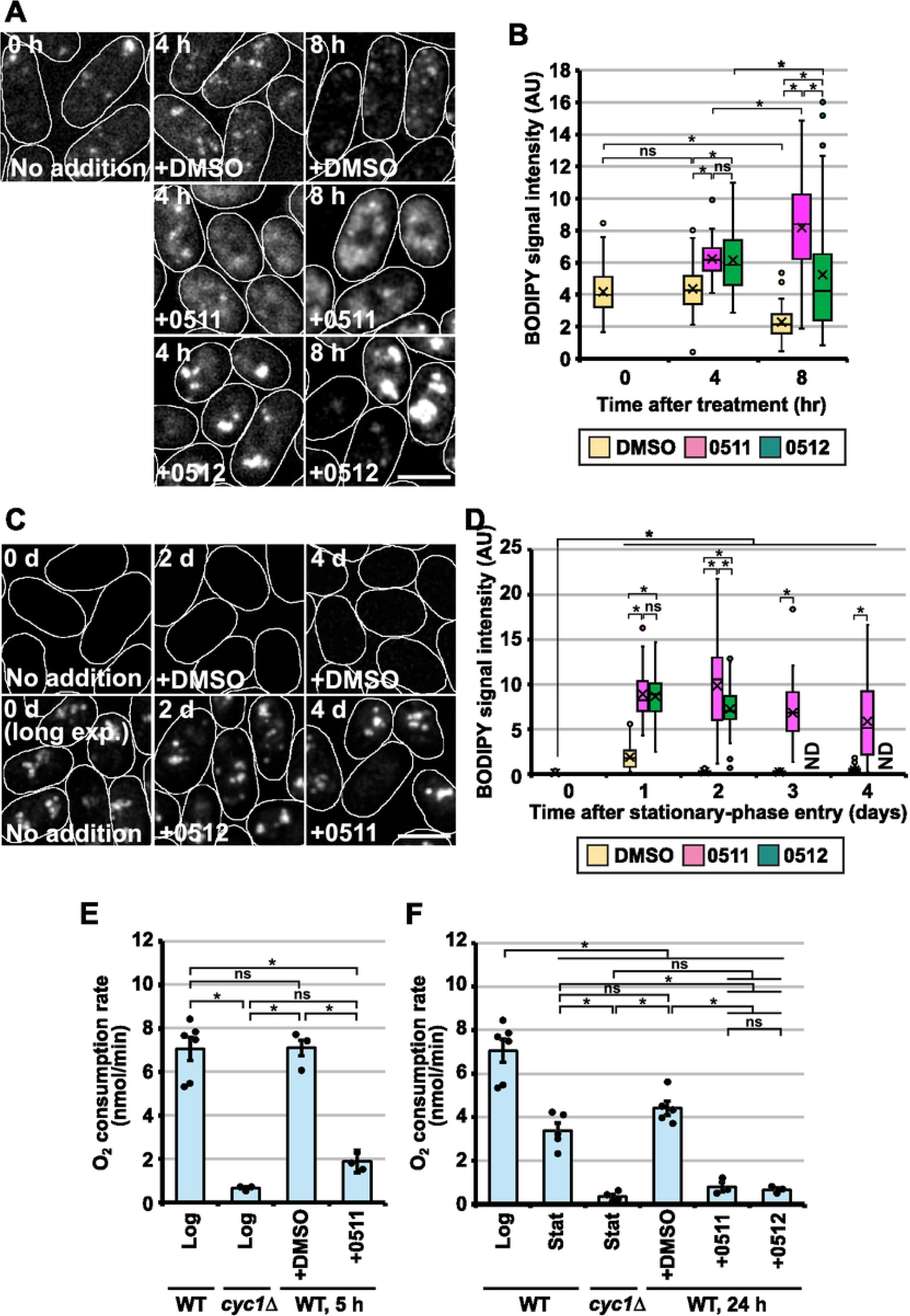
Effects of 0511 and 0512 on LDs and oxygen consumption. (A–D) BODIPY-stained LDs (A, C) and quantification of BODIPY signal intensities (B, D) in log-phase (A, B) or stationary-phase cells (C, D) treated with DMSO, 0511 or 0512. Log- or stationary-phase cells were treated with the indicated compounds (30 µM for 0511 and 0512) for the indicated periods and subsequently stained with BODIPY to visualize intracellular LDs. Treatment of log-phase cells for 4 hours does not induce cell cycle arrest, whereas treatment for 8 hours induces stationary phase-like, cell cycle arrest (see Figs 3 and 4). Images in (A) and (C) were acquired using the same exposure time, except for stationary-phase cells at day 0, which were imaged with an additional longer exposure time for visualization (long exp). White lines indicate cell outlines. Bar: 5 µm. In (B) and (D), intracellular BODIPY signal intensity was measured in more than 50 cells at each time point in one representative experiment from two (for log-phase cells) or three (for stationary-phase cells) independent experiments. In box-and-whisker plots, the horizontal line indicates the median, the box indicates IQR, whiskers show the remaining distribution within 1.5×IQR, circles outside of whiskers represent outliers, and the cross indicates the mean value. (E and F) Mitochondrial oxygen consumption rates in log-phase cells (E) and stationary-phase or stationary phase-like cells (F). Wild-type or *cyc1Δ* cells were grown at 30°C in YES medium to 5 × 10^6^ cells/mL (Log) or 1.2 × 10^8^ cells/mL (Stat). Log-phase wild-type cells were treated with 0511, 0512, or DMSO for 5 (WT, 5 h) or 24 hours (WT, 24 h). Each dot represents the oxygen consumption rate from an individual experiment, and horizontal bars represent the mean rates. Error bars: SEM. In all graphs, asterisks represent statistically significant differences (*P*<0.05). ns: not significant difference. All statistical analyses were performed using an unpaired, two-tailed Student’s *t-*test with the Bonferroni correction. Strain used for analysis: L972 and SRS6-2D.

Mitochondria play crucial roles in lipid metabolism and the survival of stationary-phase cells [50–54]. Given the reduced viability and the probable impairment of lipid metabolism caused by the compounds, we hypothesized that they might also affect mitochondrial function. To test this idea, we examined their effects on mitochondrial oxygen consumption. Log-phase cells consumed oxygen, as indicated by a decrease in oxygen concentration in the culture medium, and this consumption depended on mitochondria because it required the *cyc1^+^* gene encoding cytochrome c, an essential component for the mitochondrial electron transport chain (Fig 6E; S3C Fig). A 5-hour DMSO treatment showed no detectable effect, whereas compound treatment significantly reduced mitochondrial oxygen consumption (Fig 6E; S3C Fig), although compound treatment did not affect cell cycle progression (Fig 3B). After 24 hours, DMSO-treated cells entered stationary phase (Fig 3B) and exhibited reduced oxygen consumption, similar to non-treated cells (Fig 6F; S3D Fig). In contrast, compound-treated cells, which entered a stationary phase-like state before the cells reached stationary-phase density (Figs 3B and 4B), showed a marked reduction in mitochondrial oxygen consumption similar to stationary-phase *cyc1Δ* cells (Fig 6F; S3D Fig). These results demonstrate a substantial reduction in mitochondrial oxygen consumption in compound-treated cells.

### A lipid synthesis inhibitor cerulenin produces outcomes partly similar to those of 0511 and 0512

To further assess the effects of 0511 and 0512 on lipid metabolism, we examined whether inhibition of lipid synthesis by cerulenin, an inhibitor of fatty acid synthetase [55], results in similar outcomes. When stationary-phase cells were treated with cerulenin for 24 hours, they failed to re-enter the cell cycle by replenishing nutrients (Fig 7A). This inhibition of cell cycle resumption was dependent on the continued presence of cerulenin, as its removal allowed cells to restart the cell cycle. These results indicate that, similar to 0511 and 0512, cerulenin inhibits cell cycle re-entry of stationary-phase cells. However, unlike 0511 and 0512, cerulenin did not decrease the viability of stationary-phase cells (Fig 7B). In log-phase cells, cerulenin induced delayed cell cycle arrest similar to 0511 and 0512, but this arrest occurred more rapidly (after about one round of cell division) (Fig 7C) [56]. Cerulenin also caused rapid cell death [56], in contrast to the slower decline in cell viability caused by 0511 or 0512 (Figs 3E and 7D).

**Fig 7.**
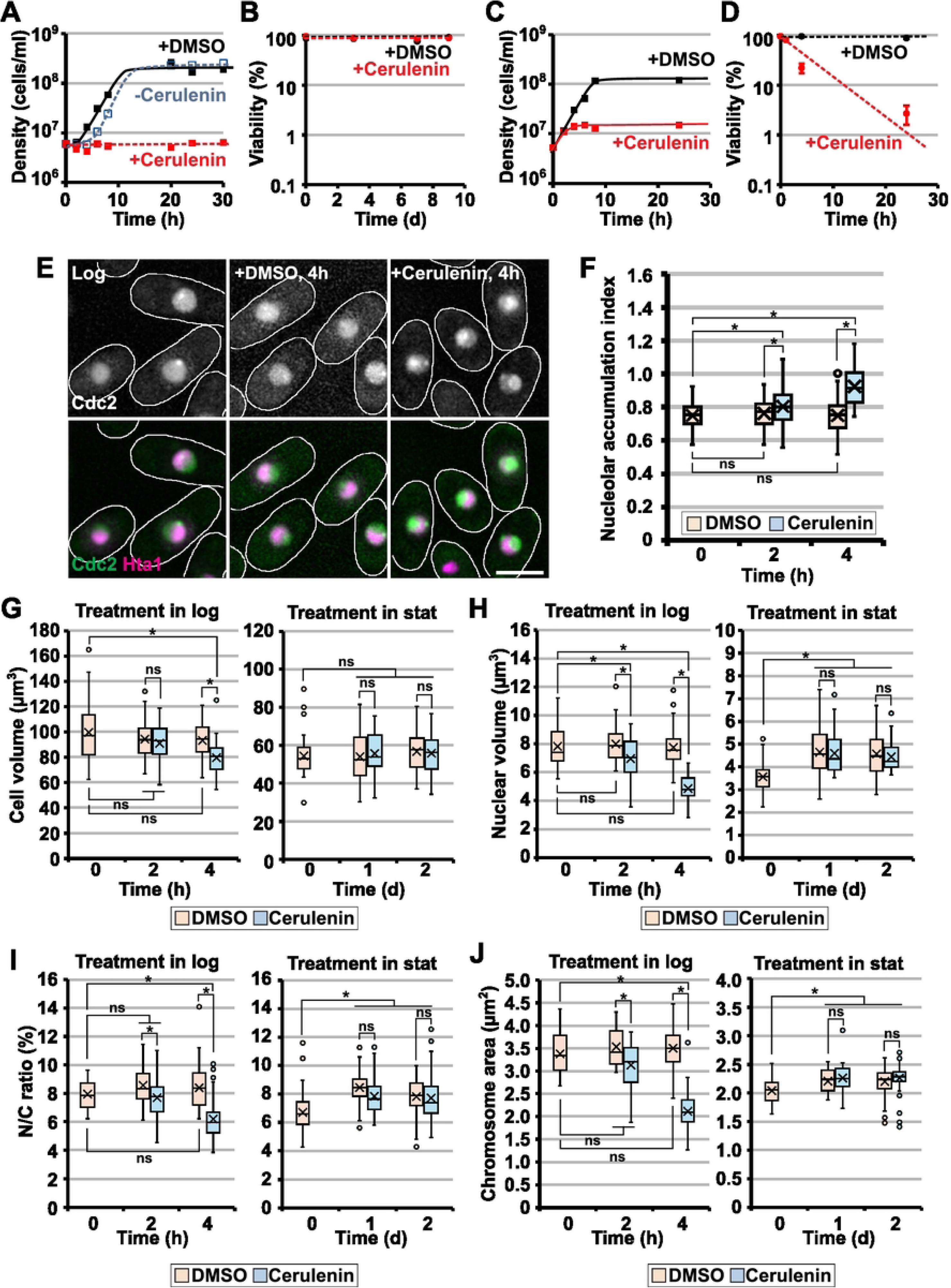
Effects of cerulenin on log-phase and stationary-phase cells. (A and B) Effects of cerulenin on cell cycle re-entry (A) and cell viability (B) in stationary phase. (C and D) Effects of cerulenin on proliferation (C) and viability (D) of log-phase cells. Cells were treated with cerulenin at 10 µg/mL. In (B) and (D), mean values from three independent experiments are shown for each time point. Error bars: SEM. (E) Intracellular Cdc2 localization in cells treated with cerulenin during log phase. White lines indicate cell outlines. Bar: 5 µm. (F) Effects of cerulenin on nucleolar accumulation index of Cdc2 in log-phase cells. (G–J) Effects of cerulenin on cell size (G), nuclear size (H), the N/C ratio (I), and chromosomal area (J) in cells treated with cerulenin. Treatment in log: chemical treatment in log phase; Treatment in stat: chemical treatment in stationary phase. In box-and-whisker plots, the horizontal line indicates the median, the box indicates IQR, whiskers show the remaining distribution within 1.5×IQR, circles outside of whiskers represent outliers, and the cross indicates the mean value. More than 50 cells were analyzed for each plot. Asterisks denote statistically significant differences (*P*<0.05). ns: not significant difference. Statistical analyses were performed using an unpaired, two-tailed Student’s *t-*test with the Bonferroni correction. All graphs and images, except those shown in (B) and (D), are from one representative experiment of two independent replicates. Strains used: L972 (A–D), SMG25-1A (E and F), and SMHT16-1 (G–J).

We next examined the effects of cerulenin on Cdc2 localization and the sizes of the cell, nucleus, and chromosomes. Cerulenin treatment during log phase induced nucleolar accumulation of Cdc2 and reduced cell size upon cell cycle arrest (Fig 7E–7G; S4A and S4B Fig), suggesting that, similar to 0511 or 0512, cerulenin induces a stationary phase-like state. Cerulenin treatment during stationary phase did not alter nucleolar accumulation of Cdc2 (data not shown) or cell size as observed with 0511 or 0512 treatment (Figs 5A and 7G, Treatment in stat; S4B Fig). Despite these similarities, cerulenin affected nuclear and chromosomal sizes in a manner markedly different from that of 0511 or 0512. In log-phase cells, cerulenin reduced nuclear and chromosomal sizes as well as the N/C ratio (Fig 7H–7J, Treatment in log; S4B Fig). Moreover, cerulenin did not suppress the DMSO-induced increase in nuclear and chromosomal sizes or the N/C ratio in stationary-phase cells (Fig 7H–7J, Treatment in stat; S4B Fig). Unlike 0511 and 0512, which increased intracellular BODIPY signal intensity, cerulenin had no effect on the dot-like BODIPY-staining pattern and caused a slight decrease or no change in signal intensity in log- or stationary-phase cells, respectively (S5A-S5D Fig). Mitochondrial oxygen consumption was also unaffected by cerulenin (S5E Fig). Taken together, these similarities and differences suggest that although 0511 and 0512 impair lipid metabolism similarly to cerulenin, their mechanisms of action probably differ.

### 0511 and 0512 reduce viability of budding yeast cells in stationary phase

We further examined whether 0511 and 0512 reduce the viability of stationary-phase cells of the budding yeast, *Saccharomyces cerevisiae,* a species evolutionarily distant from *S. pombe.* As observed in *S. pombe,* 0511 or 0512 caused a more rapid decline in the viability of stationary-phase cells compared with the DMSO control, with 0512 leading to a faster loss of viability than 0511 (Fig 8A). These findings suggest that the effects of 0511 and 0512 may be conserved between these two yeast species and that survival of quiescent yeast cells in both species may rely on similar mechanisms.

**Fig 8.**
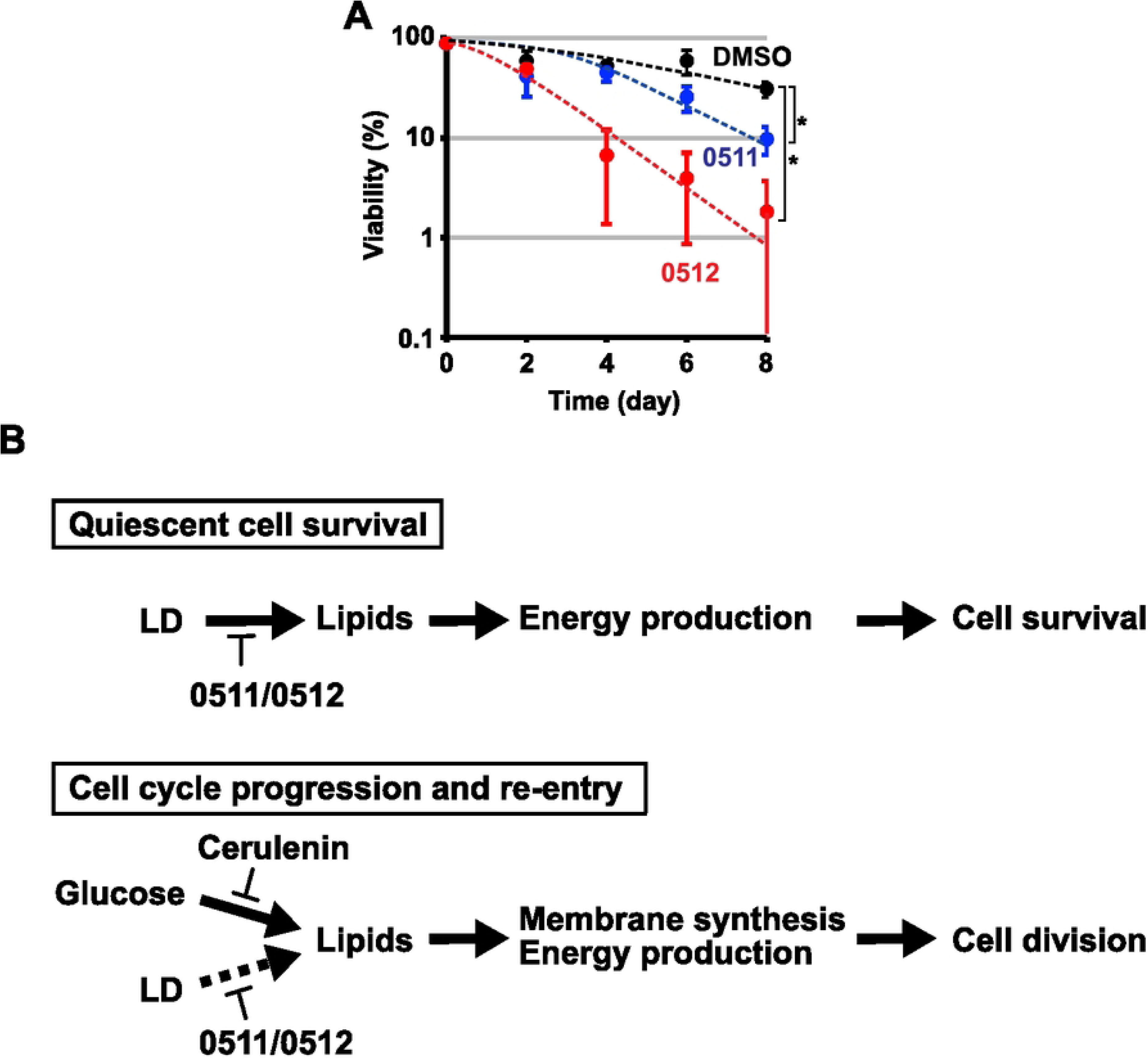
Effects of 0511 and 0512 on the viability of *S. cerevisiae* stationary-phase cells and a model for the role of lipid metabolism in *S. pombe*. (A) Effects of 0511 and 0512 on the viability of *S. cerevisiae* stationary-phase cells. *S. cerevisiae* cells were incubated at 30°C for 7 days to reach stationary phase. The cell density was then adjusted to 5 × 10^6^ cells/mL, and 0511 or 0512 was added to the culture at a final concentration of 30 µM. Cells were further incubated, and viability was assessed every 2 days. Asterisks indicate statistically significant differences compared with the DMSO control as determined by Dunnett’s test (*P<*0.05). (B) A model for the role of lipid metabolism in *S. pombe*. During stationary phase, lipids are likely supplied primarily from LDs, which are likely utilized for energy production required for survival under nutrient-poor conditions (Quiescent cell survival). 0511 and 0512 likely affect lipid mobilization from LDs, thereby reducing cell viability, whereas cerulenin, a lipid synthesis inhibitor, has negligible effects, as lipid synthesis is likely reduced in stationary phase. During proliferation and cell cycle re-entry, lipids are thought to be primarily supplied through *de novo* synthesis from glucose in the growth medium (Cell cycle progression and re-entry). Lipid supply from LDs is also important, but its contribution is smaller than that of lipid synthesis (indicated by a dotted arrow). These lipids are used for membrane synthesis and energy production required for cell division. Cerulenin inhibits cell division more rapidly than 0511 and 0512, likely because lipid synthesis contributes more to cell division than lipid mobilization from LDs. For the same reason, cerulenin inhibits cell cycle re-entry, whereas 0511 does so only after one round of resumed cell division.

## DISCUSSION

In this study, we identified novel structurally related compounds that exert unique effects on both stationary-phase and log-phase cells. In stationary-phase cells, these compounds reduced cell viability, blocked cell cycle re-entry, altered LDs, and prevented DMSO-induced enlargement of the nucleus and chromosomes. In log-phase cells, the compounds decreased viability and induced cell cycle arrest after several cell divisions, accompanied by nucleolar accumulation of Cdc2, a hallmark of stationary phase. They also inhibited mitochondrial oxygen consumption. Several of these effects overlapped with those caused by the fatty acid synthesis inhibitor, cerulenin. Similar to the compounds, cerulenin inhibited cell cycle re-entry of stationary-phase cells, induced cell cycle arrest with nucleolar accumulation of Cdc2, and reduced viability in log-phase cells. However, unlike the compounds, cerulenin did not reduce viability or suppress DMSO effects on nuclear and chromosomal size in stationary-phase cells. Cerulenin also reduced nuclear and chromosomal size in log-phase cells and had little or no effect on LDs or mitochondrial oxygen consumption, respectively. Moreover, the compounds reduced the viability of stationary-phase cells in *S. cerevisiae,* indicating that their effects may be conserved across fungal species.

The similarities between the effects of the compounds and cerulenin, together with the monoacylglycerol-like structures of the compounds, suggest that the compounds may affect lipid metabolism. The increase in intracellular BODIPY signal intensity further raises the possibility that, unlike cerulenin, the compounds affect the mobilization of lipids stored in LDs. However, the increased signal intensity should be interpreted with caution, as it may reflect altered LD composition or potential interactions between BODIPY and the compounds. If the compounds indeed perturb lipid mobilization, the differences between the effects of the compounds and cerulenin may reflect distinct cellular dependencies on lipid mobilization versus synthesis in stationary phase (Fig 8B). In stationary phase, lipid synthesis is likely suppressed due to limited nutrient availability, suggesting that cells may rely more heavily on LD-stored lipids for survival (Fig 8B, Quiescent cell survival). This difference in metabolic dependency may contribute to the reduction in cell viability in stationary-phase cells treated with the compounds, but not cerulenin. It may also explain the increase in BODIPY signal intensity, which is not observed with cerulenin. Recent studies have shown that LDs originate from the nuclear membrane [57–61] and are involved in chromosome regulation [62–65]. Interestingly, DMSO treatment resulted in changes in nuclear and chromosomal size, and the compounds modulated these changes. Although the underlying mechanism of DMSO-dependent size changes remains unclear, it is possible that DMSO also affects lipid dynamics, and the compounds interfere with this process.

In contrast, cell cycle progression and cell cycle re-entry appear to depend primarily on newly synthesized lipids [56], although stored lipids are also utilized for these processes [41,42] (Fig 8B, Cell cycle progression and re-entry). Because both lipid synthesis and mobilization are likely important for cell proliferation, inhibition of either process by the compounds or cerulenin may induce stationary phase-like cell cycle arrest and reduced cell viability as a result of lipid and energy shortages. A greater dependence on lipid synthesis during proliferation may account for the more rapid cell cycle arrest induced by cerulenin compared to the compounds, as well as a reduction in nuclear and chromosomal size observed in log-phase cells. Furthermore, LD studies in mammalian cells have suggested a close link between lipid metabolism and mitochondrial respiration [66]. In this context, the reduced mitochondrial oxygen consumption associated with compound treatment may reflect impaired lipid mobilization. Alternatively, the compounds may primarily inhibit mitochondrial functions, with secondary effects on lipid metabolism.

The precise effects and mechanisms of the compounds remain to be determined. However, given their monoacylglycerol-like structures and the similarities to cerulenin, it is tempting to speculate that the compounds interact with enzymes involved in lipid metabolism, although this remains to be experimentally tested. At the same time, the effects of the compounds on stationary-phase cells support the idea that lipid mobilization plays an important role in both cell cycle re-entry and survival during stationary phase [39–42]. Furthermore, these findings suggest that targeting lipid mobilization could represent a potential strategy for preventing the proliferation and survival of quiescent populations of pathogenic microorganisms. Indeed, *Mycobacterium tuberculosis,* a well-known bacterial pathogen that causes tuberculosis, utilizes lipids during its dormant state, and chemical compounds that target lipid metabolism have been shown to be effective against *M. tuberculosis* [67–69]. Since the compounds identified in this study do not resemble previously identified chemicals and exhibit toxicity toward evolutionarily distant yeast species, they may be useful for future studies aimed at developing novel strategies to target quiescent pathogens. In conclusion, these findings highlight the importance of lipid metabolism in survival of quiescent cells and suggest that distinct aspects of lipid regulation, such as lipid mobilization, may be critical under nutrient-limited conditions. These results provide a framework for understanding how metabolic adaptations support long-term survival in non-proliferating cells and may have broader implications for targeting quiescent or persistent cell populations.

## Acknowledgements

Screening of the compounds in this work was carried out under the project, “Recruit Innovative Ideas to Generate Original Targets Takeda”, which was organized by Takeda Pharmaceutical Company Limited. We thank Yoko Kimura for a budding yeast strain and Bas Teusink for a NeonGreen plasmid. This work was in part supported by JSPS KAKENHI (Grant-in-Aid for Scientific Research (C), Grant Number JP24K09435) to Ayumu Yamamoto.

## Supporting Information

**S1 Fig. Pairwise statistical analysis of NAI, cell and nuclear volumes, the N/C ratio, and chromosome area in log-phase (LP) cells treated with 0511 or 0512.** *P-*values were calculated using two-tailed Student’s *t-*test and adjusted by Bonferroni correction. Cells with orange background indicate comparisons with statistically significant *P-*values.

**S2 Fig. Pairwise statistical analysis of cell and nuclear volumes, the N/C ratio, and chromosome area in stationary-phase (SP) cells treated with 0511 or 0512.** *P-*values were calculated using two-tailed Student’s *t-*test and adjusted by Bonferroni correction. Cells with orange background indicate comparisons with statistically significant *P-*values.

**S3 Fig. Pairwise statistical analysis of BODIPY signal intensities and mitochondria-dependent oxygen consumption in cells treated with 0511 or 0512.** (A and B) BODIPY signal intensities in log-phase (A) or stationary-phase cells (B). (C and D) Pairwise statistical analysis of mitochondrial oxygen consumption in proliferating cells (C) and in cells in the stationary phase or stationary phase-like state (D). *P-*values were calculated using two-tailed Student’s *t-*test and adjusted by Bonferroni correction. Cells with orange background indicate pairs with significant *P-*values.

**S4 Fig. Pairwise statistical analysis of cells treated with cerulenin.** (A) Statistical analysis of nucleolar accumulation index (NAI) of Cdc2 in LP cells treated with cerulenin. (B) Statistical analysis of cell and nuclear volumes, the N/C ratio, and chromosomal area in log-phase (LP) or stationary-phase (SP) cells treated with cerulenin. *P-*values calculated using two-tailed Student’s *t-*test are shown. Values indicate *P-*values after Bonferroni correction. Cells with orange background indicate comparisons with statistically significant *P-*values.

**S5 Fig. Effects of cerulenin on LDs and mitochondria-dependent oxygen consumption.** BODIPY-stained LDs (A and C) and BODIPY signal intensities (B and D) in log-phase (A and B) or stationary-phase cells (C and D) treated with DMSO or cerulenin. Log- or stationary-phase cells were treated with DMSO or 10 µg/mL cerulenin for the indicated periods, and LDs were stained with BODIPY. In (B) and (D), The horizontal line indicates the median, the box indicates IQR, whiskers show the remaining distribution within 1.5×IQR, circles outside of whiskers represent outliers, and the cross indicates the mean value. More than 50 cells were analyzed for each plot. (E) Oxygen consumption rate in cells treated with cerulenin. Log-phase cells were treated with either DMSO or 10 µg/mL cerulenin for 4 hours. Note that 4-hour DMSO treatment did not cause cell cycle arrest, while 4-hour cerulenin treatment resulted in cell cycle arrest (see Fig. 7C). Each dot represents the oxygen consumption rate from an individual experiment, and horizontal bars represent the mean rates. Error bars: SEM. \**P*<0.05; ns: not significant difference. Statistical analyses were performed using an unpaired, two-tailed Student’s *t-*test with the Bonferroni correction. Strains used: L972.

## Notes

### Competing Interest Statement

The authors have declared no competing interest.

## References

1. Rittershaus ESC, Baek S-H, Sassetti CM. The normalcy of dormancy: common themes in microbial quiescence. Cell Host Microbe. 2013;13: 643–651. doi:10.1016/J.CHOM.2013.05.012

2. Gengenbacher M, Kaufmann SHE. *Mycobacterium tuberculosis*: success through dormancy. FEMS Microbiol Rev. 2012;36: 514–532. doi:10.1111/J.1574-6976.2012.00331.X

3. Bojsen R, Regenberg B, Folkesson A. Persistence and drug tolerance in pathogenic yeast. Curr Genet. 2017;63: 19–22. doi:10.1007/s00294-016-0613-3

4. Kloehn J, Saunders EC, O’Callaghan S, Dagley MJ, McConville MJ. Characterization of metabolically quiescent leishmania parasites in murine lesions using heavy water labeling. PLoS Pathog. 2015;11: e1004683. doi:10.1371/JOURNAL.PPAT.1004683

5. Verderosa AD, Totsika M, Fairfull-Smith KE. Bacterial biofilm eradication agents: a current review. Front Chem. 2019;7: 824. doi:10.3389/FCHEM.2019.00824

6. Sun S, Gresham D. Cellular quiescence in budding yeast. Yeast. 2021;38: 12–29. doi:10.1002/yea.3545

7. Yanagida M. Cellular quiescence: are controlling genes conserved? Trends Cell Biol. 2009;19: 705–715. doi:10.1016/J.TCB.2009.09.006

8. Herman PK. Stationary phase in yeast. Curr Opin Microbiol. 2002;5: 602–607. doi:10.1016/S1369-5274(02)00377-6

9. Gray J V, Petsko GA, Johnston GC, Ringe D, Singer RA, Werner-Washburne M. “Sleeping beauty”: quiescence in *Saccharomyces cerevisiae*. Microbiol Mol Biol Rev. 2004;68: 187–206. doi:10.1128/MMBR.68.2.187-206.2004

10. Sagot I, Laporte D. The cell biology of quiescent yeast - a diversity of individual scenarios. J Cell Sci. 2019;132: jcs213025. doi:10.1242/jcs.213025

11. De Virgilio C. The essence of yeast quiescence. FEMS Microbiol Rev. 2012;36: 306–339. doi:10.1111/j.1574-6976.2011.00287.x

12. Broach JR. Nutritional control of growth and development in yeast. Genetics. 2012;192: 73–105. doi:10.1534/genetics.111.135731

13. Hiraoka M, Kiyota Y, Kawai S, Notsu Y, Yamada K, Kurashima K, et al. CDK actively contributes to establishment of the stationary phase state in fission yeast. J Cell Sci. 2023;136: jcs260727. doi:10.1242/JCS.260727/307401/AM/CDK-ACTIVELY-CONTRIBUTES-TO-ESTABLISHMENT-OF-THE

14. Joyner RP, Tang JH, Helenius J, Dultz E, Brune C, Holt LJ, et al. A glucose-starvation response regulates the diffusion of macromolecules. Elife. 2016;5: e09376. doi:10.7554/eLife.09376

15. Narayanaswamy R, Levy M, Tsechansky M, Stovall GM, O’Connell JD, Mirrielees J, et al. Widespread reorganization of metabolic enzymes into reversible assemblies upon nutrient starvation. Proc Natl Acad Sci U S A. 2009;106: 10147–52. doi:10.1073/pnas.0812771106

16. Kelkar M, Martin SG. PKA antagonizes CLASP-dependent microtubule stabilization to re-localize Pom1 and buffer cell size upon glucose limitation. Nat Commun. 2015;6: 8445. doi:10.1038/ncomms9445

17. Leitao RM, Kellogg DR. The duration of mitosis and daughter cell size are modulated by nutrients in budding yeast. J Cell Biol. 2017;216: 3463–3470. doi:10.1083/JCB.201609114

18. Laporte D, Courtout F, Pinson B, Dompierre J, Salin B, Brocard L, et al. A stable microtubule array drives fission yeast polarity reestablishment upon quiescence exit. J Cell Biol. 2015;210: 99–113. doi:10.1083/jcb.201502025

19. Laporte D, Courtout F, Salin B, Ceschin J, Sagot I. An array of nuclear microtubules reorganizes the budding yeast nucleus during quiescence. J Cell Biol. 2013;203: 585–594. doi:10.1083/JCB.201306075

20. Sagot I, Pinson B, Salin B, Daignan-Fornier B. Actin bodies in yeast quiescent cells: an immediately available actin reserve? Chang F, editor. Mol Biol Cell. 2006;17: 4645–4655. doi:10.1091/mbc.e06-04-0282

21. Buchan JR, Parker R. Eukaryotic stress granules: the ins and outs of translation. Mol Cell. 2009;36: 932–941. doi:10.1016/J.MOLCEL.2009.11.020/ATTACHMENT/5AC4E99F-526A-44CA-9DB6-1A8D932D0789/MMC1.PDF

22. Buchan JR, Muhlrad D, Parker R. P bodies promote stress granule assembly in Saccharomyces cerevisiae. J Cell Biol. 2008;183: 441–455. doi:10.1083/JCB.200807043

23. Nilsson D, Sunnerhagen P. Cellular stress induces cytoplasmic RNA granules in fission yeast. RNA. 2011;17: 120–133. doi:10.1261/RNA.2268111

24. Shah KH, Zhang B, Ramachandran V, Herman PK. Processing body and stress granule assembly occur by independent and differentially regulated pathways in *Saccharomyces cerevisiae*. Genetics. 2013;193: 109–123. doi:10.1534/GENETICS.112.146993/-/DC1

25. Laporte D, Salin B, Daignan-Fornier B, Sagot I. Reversible cytoplasmic localization of the proteasome in quiescent yeast cells. J Cell Biol. 2008;181: 737–45. doi:10.1083/jcb.200711154

26. Heimlicher MB, Bächler M, Liu M, Ibeneche-Nnewihe C, Florin EL, Hoenger A, et al. Reversible solidification of fission yeast cytoplasm after prolonged nutrient starvation. J Cell Sci. 2019;132: jcs231688. doi:10.1242/jcs.231688

27. Swygert SG, Kim S, Wu X, Fu T, Hsieh TH, Rando OJ, et al. Condensin-dependent chromatin compaction represses transcription globally during quiescence. Mol Cell. 2019;73: 533–546. doi:10.1016/J.MOLCEL.2018.11.020

28. Schäfer G, McEvoy CRE, Patterton H-G. The *Saccharomyces cerevisiae* linker histone Hho1p is essential for chromatin compaction in stationary phase and is displaced by transcription. Proc Natl Acad Sci U S A. 2008;105: 14838–43. doi:10.1073/pnas.0806337105

29. Laporte D, Courtout F, Tollis S, Sagot I. Quiescent *Saccharomyces cerevisiae* forms telomere hyperclusters at the nuclear membrane vicinity through a multifaceted mechanism involving Esc1, the Sir complex, and chromatin condensation. Bloom KS, editor. Mol Biol Cell. 2016;27: 1875–1884. doi:10.1091/mbc.e16-01-0069

30. Moreno-Torres M, Jaquenoud M, De Virgilio C. TORC1 controls G1–S cell cycle transition in yeast via Mpk1 and the greatwall kinase pathway. Nat Commun. 2015;6: 8256. doi:10.1038/ncomms9256

31. Simanis V, Nurse P. The cell cycle control gene *cdc2^+^* of fission yeast encodes a protein kinase potentially regulated by phosphorylation. Cell. 1986;45: 261–268. doi:10.1016/0092-8674(86)90390-9

32. Werner-Washburne M, Braun E, Johnston GC, Singer RA. Stationary phase in the yeast *Saccharomyces cerevisiae*. Microbiol Rev. 1993;57: 383–401. Available: http://www.ncbi.nlm.nih.gov/pubmed/8393130

33. Zinzalla V, Graziola M, Mastriani A, Vanoni M, Alberghina L. Rapamycin-mediated G1 arrest involves regulation of the Cdk inhibitor Sic1 in *Saccharomyces cerevisiae*. Mol Microbiol. 2007;63: 1482–1494. doi:10.1111/j.1365-2958.2007.05599.x

34. Oelkers P, Cromley D, Padamsee M, Billheimer JT, Sturley SL. The DGA1 gene determines a second triglyceride synthetic pathway in yeast. J Biol Chem. 2002;277: 8877–8881. doi:10.1074/jbc.M111646200

35. Handee W, Li X, Hall KW, Deng X, Li P, Benning C, et al. An energy-independent pro-longevity function of triacylglycerol in yeast. PLoS Genet. 2016;12: e1005878. doi:10.1371/JOURNAL.PGEN.1005878

36. Fernández-Murray JP, McMaster CR. Lipid synthesis and membrane contact sites: A crossroads for cellular physiology. J Lipid Res. 2016;57: 1789–1805. doi:10.1194/JLR.R070920/ASSET/FB40761F-467C-4F07-BE03-F24ED2E7323B/MAIN.ASSETS/GR4.JPG

37. Pascual F, Soto-Cardalda A, Carman GM. PAH1-encoded phosphatidate phosphatase plays a role in the growth phase- and inositol-mediated regulation of lipid synthesis in *Saccharomyces cerevisiae*. J Biol Chem. 2013;288: 35781–35792. doi:10.1074/JBC.M113.525766/ASSET/02196990-ED5A-47EF-B282-1F4F3C423E61/MAIN.ASSETS/GR9.JPG

38. Han GS, Wu WI, Carman GM. The *Saccharomyces cerevisiae* lipin homolog is a Mg^2+^-dependent phosphatidate phosphatase enzyme. J Biol Chem. 2006;281: 9210–9218. doi:10.1074/JBC.M600425200/ASSET/759FE8D3-8323-4FF0-9AAB-384C62592531/MAIN.ASSETS/GR9.JPG

39. Park Y, Han GS, Mileykovskaya E, Garrett TA, Carman GM. Altered lipid synthesis by lack of yeast Pah1 phosphatidate phosphatase reduces chronological life span. J Biol Chem. 2015;290: 25382–25394. doi:10.1074/JBC.M115.680314/ASSET/CBDB1803-48EE-41EC-B0B8-1AC1D3BCBBE3/MAIN.ASSETS/GR14.JPG

40. Seo AY, Lau PW, Feliciano D, Sengupta P, Le Gros MA, Cinquin B, et al. AMPK and vacuole-associated Atg14p orchestrate μ-lipophagy for energy production and long-term survival under glucose starvation. Elife. 2017;6: e21690. doi:10.7554/ELIFE.21690

41. Kurat CF, Wolinski H, Petschnigg J, Kaluarachchi S, Andrews B, Natter K, et al. Cdk1/Cdc28-dependent activation of the major triacylglycerol lipase Tgl4 in yeast links lipolysis to cell-cycle progression. Mol Cell. 2009;33: 53–63. doi:10.1016/j.molcel.2008.12.019

42. Kurat CF, Natter K, Petschnigg J, Wolinski H, Scheuringer K, Scholz H, et al. Obese yeast: triglyceride lipolysis is functionally conserved from mammals to yeast. J Biol Chem. 2006;281: 491–500. doi:10.1074/JBC.M508414200

43. Moreno S, Klar A, Nurse P. Molecular genetic analysis of fission yeast *Schizosaccharomyces pombe*. Methods Enzymol. 1991;194: 795–823. doi:10.1016/0076-6879(91)94059-L

44. Nambu M, Kishikawa A, Yamada T, Ichikawa K, Kira Y, Itabashi Y, et al. Direct evaluation of cohesin-mediated sister kinetochore associations at meiosis I in fission yeast. J Cell Sci. 2022;135: jcs259102. doi:10.1242/JCS.259102/273595/AM/DIRECT-EVALUATION-OF-COHESIN-MEDIATED-SISTER

45. Matsuhara H, Yamamoto A. Autophagy is required for efficient meiosis progression and proper meiotic chromosome segregation in fission yeast. Genes to Cells. 2016;21: 65–87. doi:10.1111/gtc.12320

46. Jacobus AP, Gross J. Optimal Cloning of PCR Fragments by Homologous Recombination in Escherichia coli. Korolev S, editor. PLoS One. 2015;10: e0119221. doi:10.1371/journal.pone.0119221

47. Schneider CA, Rasband WS, Eliceiri KW. NIH Image to ImageJ: 25 years of image analysis. Nat Methods. 2012;9: 671–675. doi:10.1038/nmeth.2089

48. Kume K, Cantwell H, Neumann FR, Jones AW, Snijders AP, Nurse P. A systematic genomic screen implicates nucleocytoplasmic transport and membrane growth in nuclear size control. PLoS Genet. 2017;13: e1006767. doi:10.1371/JOURNAL.PGEN.1006767

49. Meyers A, del Rio ZP, Beaver RA, Morris RM, Weiskittel TM, Alshibli AK, et al. Lipid droplets form from distinct regions of the cell in the fission yeast *Schizosaccharomyces pombe*. Traffic. 2016;17: 657–669. doi:10.1111/tra.12394

50. Pan Y, Schroeder EA, Ocampo A, Barrientos A, Shadel GS. Regulation of yeast chronological life span by TORC1 via adaptive mitochondrial ROS signaling. Cell Metab. 2011;13: 668–678. doi:10.1016/J.CMET.2011.03.018

51. Zuin A, Gabrielli N, Calvo IA, García-Santamarina S, Hoe KL, Kim DU, et al. Mitochondrial dysfunction increases oxidative stress and decreases chronological life span in fission yeast. PLoS One. 2008;3: e2842. doi:10.1371/JOURNAL.PONE.0002842

52. Ocampo A, Liu J, Schroeder EA, Shadel GS, Barrientos A. Mitochondrial respiratory thresholds regulate yeast chronological life span and its extension by caloric restriction. Cell Metab. 2012;16: 55–67. doi:10.1016/J.CMET.2012.05.013

53. Bonawitz ND, Chatenay-Lapointe M, Pan Y, Shadel GS. Reduced TOR signaling extends chronological life span via increased respiration and upregulation of mitochondrial gene expression. Cell Metab. 2007;5: 265–277. doi:10.1016/J.CMET.2007.02.009

54. Bonawitz ND, Rodeheffer MS, Shadel GS. Defective mitochondrial gene expression results in reactive oxygen species-mediated inhibition of respiration and reduction of yeast life span. Mol Cell Biol. 2006;26: 4818–4829. doi:10.1128/MCB.02360-05

55. Funabashi H, Kawaguchi A, Tomoda H, Ömura S, Okuda S, Iwasaki S. Binding site of cerulenin in fatty acid synthetase. The Journal of Biochemistry. 1989;105: 751–755. doi:10.1093/OXFORDJOURNALS.JBCHEM.A122739

56. Saitoh S, Takahashi K, Nabeshima K, Yamashita Y, Nakaseko Y, Hirata A, et al. Aberrant mitosis in fission yeast mutants defective in fatty acid synthetase and acetyl CoA carboxylase. J Cell Biol. 1996;134: 949–961. doi:10.1083/JCB.134.4.949

57. Romanauska A, Köhler A. The inner nuclear membrane Is a metabolically active territory that generates nuclear lipid droplets. Cell. 2018;174: 700–715.e18. doi:10.1016/J.CELL.2018.05.047

58. Cartwright BR, Binns DD, Hilton CL, Han S, Gao Q, Goodman JM. Seipin performs dissectible functions in promoting lipid droplet biogenesis and regulating droplet morphology. Mol Biol Cell. 2015;26: 726–739. doi:10.1091/MBC.E14-08-1303/MC-E14-08-1303-S13.MOV

59. Ohsaki Y, Kawai T, Yoshikawa Y, Cheng J, Jokitalo E, Fujimoto T. PML isoform II plays a critical role in nuclear lipid droplet formation. J Cell Biol. 2016;212: 29–38. doi:10.1083/JCB.201507122

60. Wolinski H, Hofbauer HF, Hellauer K, Cristobal-Sarramian A, Kolb D, Radulovic M, et al. Seipin is involved in the regulation of phosphatidic acid metabolism at a subdomain of the nuclear envelope in yeast. Biochim Biophys Acta. 2015;1851: 1450–1464. doi:10.1016/J.BBALIP.2015.08.003

61. Sołtysik K, Ohsaki Y, Tatematsu T, Cheng J, Maeda A, Morita SY, et al. Nuclear lipid droplets form in the inner nuclear membrane in a seipin-independent manner. J Cell Biol. 2020;220: e202005026. doi:10.1083/JCB.202005026/211592

62. Welte MA. Expanding roles for lipid droplets. Curr Biol. 2015;25: R470–R481. doi:10.1016/J.CUB.2015.04.004/ASSET/86CD0020-FAFB-4E22-AAC9-E58C81292A6B/MAIN.ASSETS/GR5.JPG

63. Welte MA, Gould AP. Lipid droplet functions beyond energy storage. Biochim Biophys Acta. 2017;1862: 1260–1272. doi:10.1016/J.BBALIP.2017.07.006

64. Gao Q, Goodman JM. The lipid droplet-a well-connected organelle. Front Cell Dev Biol. 2015;3: 154226. doi:10.3389/FCELL.2015.00049/XML

65. Henne WM, Reese ML, Goodman JM. The assembly of lipid droplets and their roles in challenged cells. EMBO J. 2018;37: e98947. doi:10.15252/EMBJ.201898947/ASSET/AFAD48E2-AFE4-4EF5-9969-1C161F3490E3/ASSETS/GRAPHIC/EMBJ201898947-FIG-0003-M.PNG

66. Varghese M, Kimler VA, Ghazi FR, Rathore GK, Perkins GA, Ellisman MH, et al. Adipocyte lipolysis affects Perilipin 5 and cristae organization at the cardiac lipid droplet-mitochondrial interface. Sci Rep. 2019;9: 1–12. doi:10.1038/s41598-019-41329-4

67. Sturm A, Sun P, Avila-Pacheco J, Clatworthy AE, Bloom-Ackermann Z, Wuo MG, et al. Genetic factors affecting storage and utilization of lipids during dormancy in *Mycobacterium tuberculosis*. mBio. 2024;15: e03208–23. doi:10.1128/MBIO.03208-23/SUPPL_FILE/MBIO.03208-23-S0002.XLSX

68. Babin BM, Keller LJ, Pinto Y, Li VL, Eneim AS, Vance SE, et al. Identification of covalent inhibitors that disrupt M. tuberculosis growth by targeting multiple serine hydrolases involved in lipid metabolism. Cell Chem Biol. 2022;29: 897–909.e7. doi:10.1016/J.CHEMBIOL.2021.08.013

69. Kim H, Shin SJ. Revolutionizing control strategies against *Mycobacterium tuberculosis* infection through selected targeting of lipid metabolism. Cellular and Molecular Life Sciences. 2023;80: 1–20. doi:10.1007/S00018-023-04914-5

